# Exploring the GM-CSF Histidine Triad as a Modulator of Structure, Molecular Motion, and Ligand Binding

**DOI:** 10.64898/2026.01.20.700583

**Authors:** Jennifer Y. Cui, Iz Varghese, Anna S. Bock, Mariana Floody, Fuming Zhang, Brenda M. Rubenstein, George P. Lisi

## Abstract

Granulocyte macrophage-colony stimulating factor (GM-CSF) is a cytokine that plays a role in immune modulation. Its expression is associated with a multitude of different effects ranging from harmful, as in diseases such as rheumatoid arthritis and multiple sclerosis, to beneficial, as in the case of mitigation of diabetes type I and neutropenia. However, there is a large gap in knowledge explaining how GM-CSF toggles its structure for such physiological and pathological interactions. Our work describes an ongoing attempt to address this gap by focusing on a clustered histidine triad within *α*-helices near the N-terminus, which prior studies have suggested play a role in binding ligands at an acidic pH. While GM-CSF is known to be highly flexible at a more acidic pH, several properties of its histidine triad remain unclear at the physiological pH at which GM-CSF would encounter its binding partners. We describe an effort to characterize the role of the GM-CSF histidines under physiological pH, specifically to determine if these histidines are key to GM-CSF structural integrity, and whether individual histidine residues modulate binding as they do at a lower pH. Our findings reveal that, while the histidine residues have an impact on GM-CSF structure, flexibility, and stability, they alone do not modulate the affinity for ligands at neutral pH. These data provide an initial explanation for the pleiotropic functions and interactions of GM-CSF within a biophysical context.

**Graphical Abstract:** 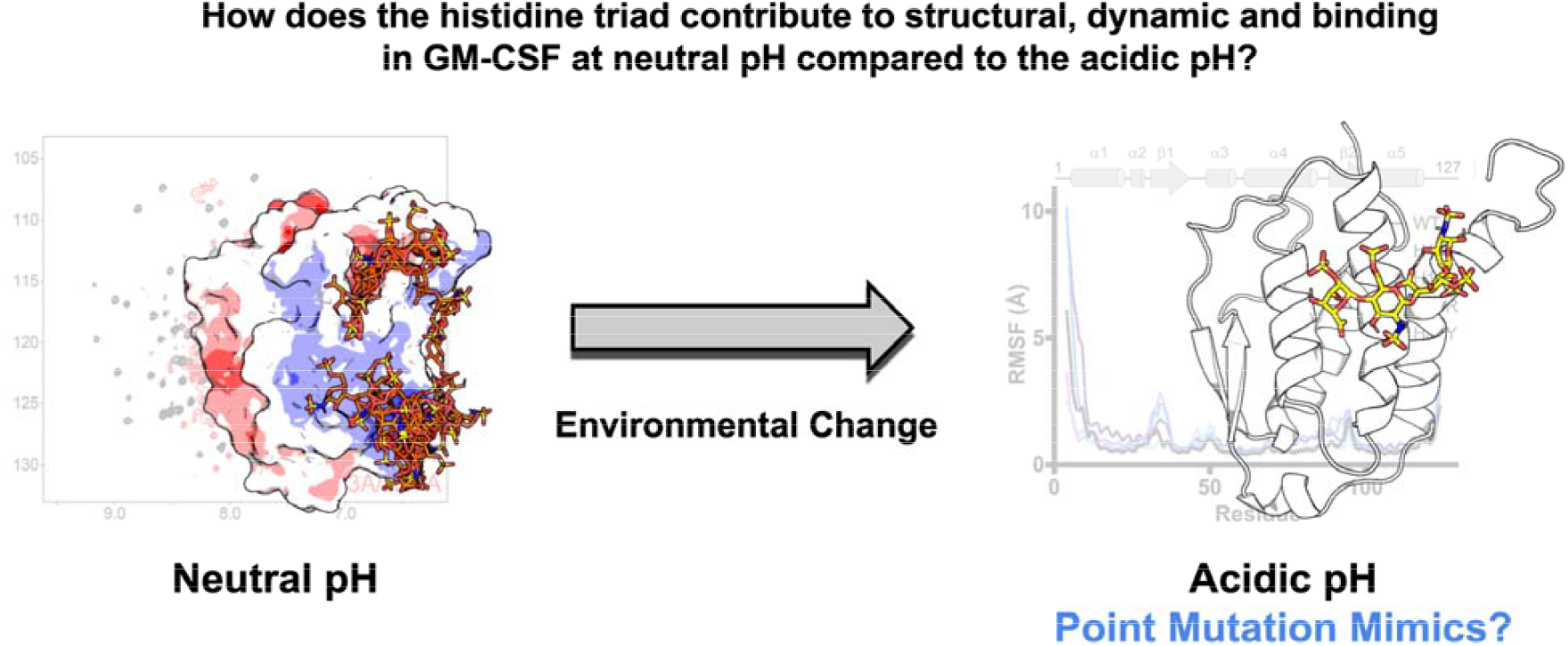

## Introduction

Cytokines are small signaling proteins that regulate immune responses under physiological and pathological conditions. The functional versatility of cytokines positions them as key mediators of cellular communication, orchestrating processes such as inflammation, tissue repair, and immune defense. Within this protein family, granulocyte macrophage-colony stimulating factor (GM-CSF) is a particularly intriguing example due to its pleiotropic roles in health and disease.

GM-CSF and other colony stimulating factors were originally shown to promote proliferation of myeloid progenitor cells. GM-CSF has a more limited role during steady-state myelopoiesis^1, 2^, confined to the development and maintenance of alveolar macrophages and non-lymphoid dendritic cells^1^. The immunomodulatory^3^ and pro-inflammatory signaling functions^4-6^ of GM-CSF have been shown to aggravate conditions such as rheumatoid arthritis and multiple sclerosis^7, 8^. Contrasting with its role in inflammation is the ability of GM-CSF to ameliorate diseases such as type-I diabetes^9^ and its clinical use to combat neutropenia^10^. GM-CSF expression levels have also been implicated in protection against *Mycobacterium tuberculosis* infection, an increase in phagocytic capacity of alveolar macrophages ^11^, and dose-dependent protection against HIV replication^12^. This functional diversity strongly suggests additional axes of regulation for GM-CSF, beyond its localization.

A substantial knowledge gap exists in our understanding of the ability of cytokines, including GM-CSF, to engage in non-overlapping cellular interactions despite small and seemingly unremarkable structures. A growing body of evidence suggests that cytokines toggle their structures in response to various microenvironments^13^; thus, leveraging intrinsic structural plasticity could be the focus of future bioengineering efforts to spatiotemporally control these molecular interactions. By manipulating specific amino acids within cytokine structures, we can establish the most critical regions for tuning conformational dynamics, leading to new strategies for regulating cytokine activity in immune responses. However, molecular insight into the biophysical characteristics that define the binding properties and mechanisms of many cytokines is lacking.

Our work describes an ongoing attempt to understand this interplay in GM-CSF, where prior studies suggested that a histidine triad located on alpha-helices near the N-terminus plays a role in binding ligands at an acidic pH^14^. Further research from our group characterized heparin-dependent structural changes in GM-CSF using NMR, and the extent to which these changes were driven by both the length of the heparin chain and the solution pH^15^. As in prior work, we hypothesized these effects to be linked to the histidine triad within GM-CSF, though we never explored contributions of the histidines directly.

At acidic pH (≤5.5), GM-CSF is known to be highly flexible, but several properties of its histidine triad remain unclear at the physiological pH at which GM-CSF would encounter binding partners. The goal of this current work is to characterize the role of the uniquely placed GM-CSF histidines as a structural switch via site-directed mutagenesis under physiological pH. We aimed to understand if disruption of specific histidine residues could mimic the pH-dependent effects observed earlier and to evaluate the overall role of each histidine in maintaining GM-CSF structural integrity. Our findings reveal that single-point mutations of the GM-CSF histidine residues have a significant impact on its structure, flexibility, and stability. Despite these biophysical impacts, mutations have limited effects on the affinity of GM-CSF for several ligands. These data, therefore, present a picture in which the histidine triad plays a significant role in the maintenance of local N-terminal structure, but a smaller part in ligand binding within a positively charged face of GM-CSF.

## Materials and Methods

### Protein expression and purification

Plasmid DNA containing GM-CSF with an N-terminal His_6_ tag was cloned into a pET-15b vector and transformed into BL21(DE3) cells. GM-CSF used for biophysical experiments was expressed in LB medium at 37 °C. Isotopically enriched GM-CSF for NMR studies was expressed at 37 °C in M9 minimal medium supplemented with CaCl_2_, MgSO_4_, and MEM vitamins, with ^15^NH_4_Cl and ^13^C_6_H_12_O_6_ as the sole nitrogen and carbon sources, respectively. For isotopic expression, 15 mL cultures of GM-CSF were first grown overnight in LB medium. The following morning, cloudy suspensions were collected by centrifugation and resuspended in the final M9 growth medium. These cultures were grown at 37 °C to an OD_600_ of 0.8-1.0 before induction with 1 mM isopropyl β-D-1-thiogalactopyranoside (IPTG). T h e cells were harvested after 5 hours of additional shaking at 37 °C and resuspended in a denaturing lysis buffer containing 10 mM Tris-HCl, 100 mM sodium phosphate, and 6 M guanidine hydrochloride (GuHCl) at pH 8.0. The cells were lysed by sonication and cell debris was removed by centrifugation.

The resulting supernatant was incubated with 10 mL of Ni-NTA agarose beads for 30 min at room temperature on a nutator before the Ni-NTA slurry was packed into a gravity column. The column was washed with the initial lysis buffer, followed by a 100 mL gradient of the same buffer without GuHCl. Elution of GM-CSF in its denatured state was performed with 1 column volume of a buffer containing 10 mM Tris-HCl, 100 mM sodium phosphate, and 250 mM imidazole at pH 8.0. GM-CSF was refolded by dilution via dropwise addition of the 10 mL eluent into 100 mL of a refolding buffer containing 10 mM Tris-HCl, 100 mM sodium phosphate, and 750 mM arginine at pH 8.0. The refolded protein was dialyzed exhaustively against a storage buffer containing 2 mM sodium phosphate at pH 7.4. GM-CSF was concentrated to 200 µM with an Amicon centrifugal device and stored at -20 °C.

### NMR spectroscopy

NMR samples were prepared by dialyzing 200 µM GM-CSF against a buffer of 20 mM HEPES and 1 mM EDTA at pH 7.4. NMR experiments were performed on Bruker Avance NEO 600 MHz and Bruker Avance III HD 850 MHz spectrometers at 25 °C and the data were processed with NMRPipe^16^ and analyzed with NMRFAM-Sparky^17^. NMR assignments of wild-type (WT) GM-CSF were initially determined with triple-resonance experiments at pH 7.4 and confirmed by BMRB entry 15531. Assignments of GM-CSF variants were mapped using the WT assignments and BMRB entry 15531 or confirmed by additional HNCO, HNCA, HNCACB, and HN(CO)CACB experiments. NMR chemical shift perturbation (CSP) analyses^18^ were carried out using ^1^H^15^N HSQC pulse programs^19^ with the ^1^H and ^15^N carrier frequencies set to the water resonance and 120 ppm, respectively. NMR chemical shift perturbations were calculated as:

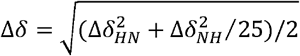

CSP significance was determined by a 10% trimmed mean of all data sets. Ligand titration experiments were performed by adding either heparin or ATP (New England BioLabs P0756S) in small concentrations to the sample until saturation or no additional perturbations were observed. Small uniform heparins with exactly 6 repeating disaccharide units (degree of polymerization 6, dp6) were used. Although in a medical context, dp6 would be considered exceedingly small, this size was selected to assess binding such that one molecule of heparin could not bind multiple sites on GM-CSF. Apparent *K*_d_ was calculated by tracking combined ^1^H^15^N CSPs of isolated peaks at each concentration and subsequent curve fitting of the points using the GraphPad Prism 10.0 *K*_d_ tool (GraphPad Software).

Longitudinal^20^ and transverse^21^ relaxation rates were determined from peak intensities of each amide resonance at multiple delay points. *T*_1_ delays of 0, 20, 60, 100, 200, 600, 800, 1200, 1500, and 2000 ms and *T*_2_ delays of 0, 16.9, 33.9, 67.8, 136.0, 169.0, and 203.0 ms were used. Peak intensities were quantified in Sparky and the resulting decay profiles were analyzed in Sparky with errors determined from the fitted parameters. Uncertainties in these rates were determined from replicate spectra with duplicate relaxation delays of 20 (x2), 60 (x2), 100, 200, 600 (x2), 800, 1200, 1500, and 2000 ms for *T*_1_ and 16.9, 33.9 (x2), 67.8, 136.0 (x2), 169.0, and 203.0 (x2) ms for *T*_2_. The heteronuclear cross-relaxation rate^21^ (NOE) was obtained by interleaving pulse sequences with and without proton saturation and calculated from the ratio of peak heights from these experiments (I_sat_/I_ref_.). All relaxation experiments were carried out in a temperature-compensated interleaved manner, processed with in-house scripts, and analyzed in GraphPad Prism 10.0.

### Circular dichroism spectroscopy

Circular dichroism (CD) spectra were measured on a Jasco J-815 spectropolarimeter in a 0.2 cm quartz cuvette with 10 µM GM-CSF in a buffer of 20 mM sodium phosphate at pH 7.4. Variable temperature measurements were collected at 208 or 220 nm over a temperature range of 25 – 90 °C, sampling every 1.5 °C at a rate of 1.5 °C/min. Unfolding profiles were fit and analyzed in GraphPad Prism 10.0 using the following equation.

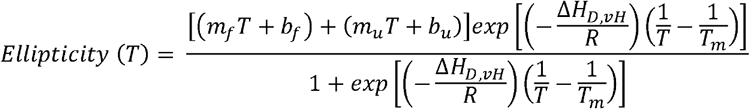

where *m*_f_ and *b*_f_ represent the slope and y-intercepts, respectively of the folded (low temperature) region of the curve and *m*_u_ and *b*_u_ represent the same values in the unfolded (high temperature) region.

### Molecular dynamics simulations

Molecular dynamics (MD) simulations based on the X-ray structure of GM-CSF (PDB: 2GMF) were performed using the OpenMM library (version 8.1.2)^22^. The AMBER ff14SB force field was used for the protein with the TIP3P water model^23^. In each simulated system, the number of Na^+^ and Cl^-^ ions was adjusted using PDBFixer to provide a physiologically-relevant ionic strength of 0.15 M^22^. Simulations were performed using an NPT ensemble with a Langevin Integrator and a Monte Carlo barostat to maintain a constant temperature and pressure of 310 K and 1 atm. Long-range electrostatic interactions were calculated using the particle mesh Ewald method^24^, with a cutoff of 1.0 nm, and an Ewald error tolerance of 0.0005. The equations of motion were integrated using a Langevin Middle Integrator, with a friction coefficient of 1.0 ps□^1^ and a time step of 2 fs. Each system was equilibrated for 100 ns, followed by a production run of 250 ns. Simulation images were created using PyMOL^25^, and plots were prepared using GraphPad Prism 10.0. Solvent Accessible Surface Area (SASA) for the entire protein was calculated in VMD^26^ using a probe radius of 1.4 Å. Electrostatic surfaces were constructed using the APBS Electrostatic plugin^27^ on PyMOL.

### Protein structure prediction

Protein fold similarity searches and predictions were carried out using the AlphaFold 3 server^28^. Prediction accuracy was assessed using template modeling score (TM-score), and local distance difference test (pLDDT). Results were visualized using PyMOL^25^.

## Results

### The GM-CSF histidine triad is a critical anchor point of structural stability at neutral pH

Histidine residues are integral to the structural and functional properties of many proteins, often facilitating ligand binding and catalytic activity. Clusters of histidine residues have been shown to facilitate phase separation^29, 30^, cellular pH sensing^31, 32^, and metal binding^33, 34^. In the context of GM-CSF, three histidine residues are arranged in a manner that appear to modulate its N-terminal *α*-helix (**Figure 1A-C**), serving as a basis for conformational changes via intrinsic dynamics or the binding of negatively charged ligands^14, 15, 35^. Although the histidines are close in three-dimensional space to one another, the level of solvent exposure at each site differs significantly (**Figure 1B**).

**Figure 1.**
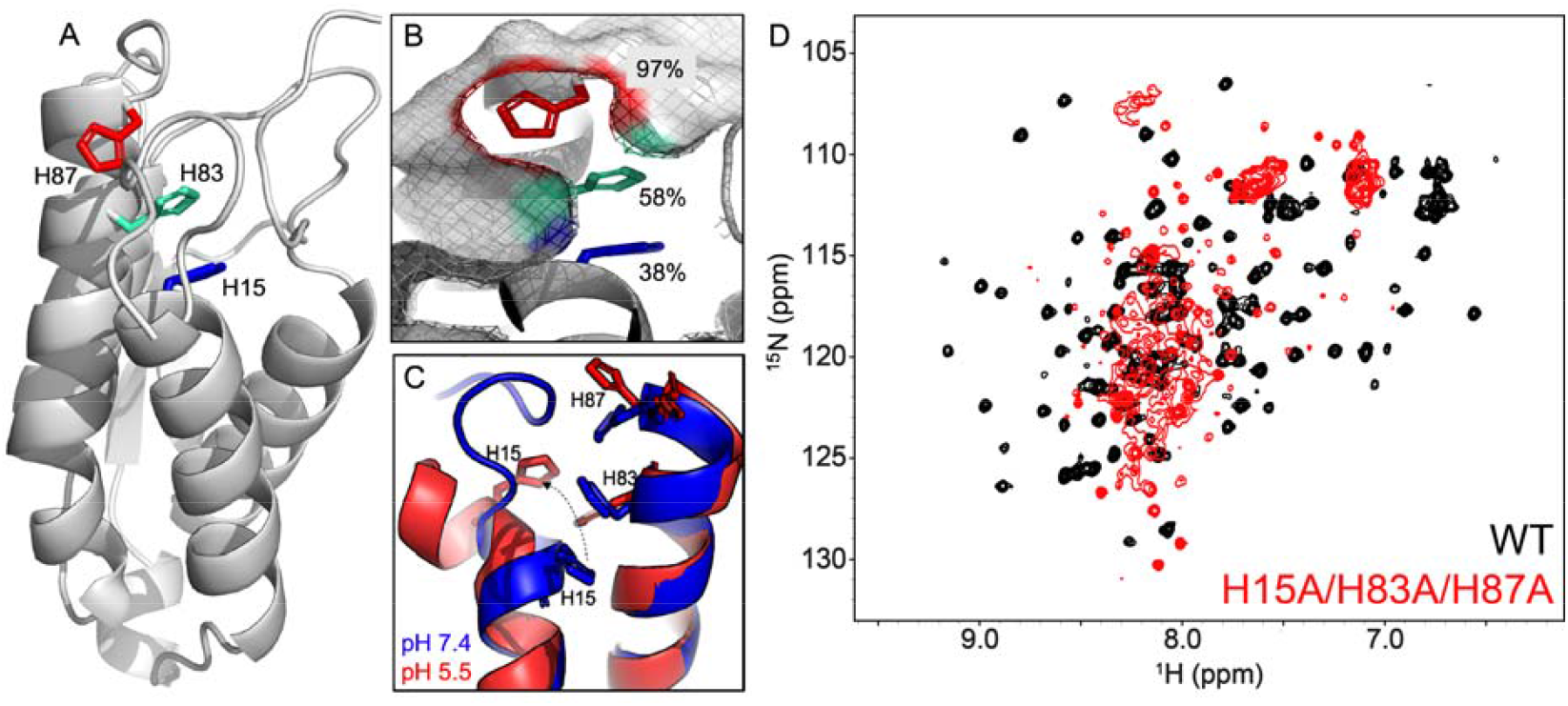
Properties of the GM-CSF histidine triad. **(A)** Structure of the GM-CSF protein (PDB: 2GMF) showing the histidine residues as sticks. **(B)** Stacking of histidine residues at the N-terminal helix of GM-CSF. The percent solvent exposure is indicated at each site. **(C)** pH-dependent conformational change of the GM-CSF helix, disrupting the stacking of histidines. **(D)** The ^1^H^15^N HSQC NMR spectrum of H15A/H83A/H87A GM-CSF (red) shows a narrow resonance dispersion indicative of an aggregated or unstructured protein. The ^1^H^15^N HSQC NMR spectrum of folded WT GM-CSF is shown in black, for reference.

To investigate the contribution of the entire histidine triad to GM-CSF’s structural integrity, we created a His-to-Ala triple mutant that produced a highly unstable GM-CSF protein. Although the protein remained soluble, it was largely unfolded. The NMR spectrum of this variant displayed overlapping and unresolved resonances clustered in the center of the spectrum (**Figure 1D**), reflecting a loss of structural integrity typical of misfolded or disordered proteins. Though the GM-CSF histidines confer flexibility to the protein when protonated, it was somewhat surprising that His-to-Ala mutations would so strongly disrupt its structure at neutral pH.

### NMR spectra of GM-CSF variants highlight His solvent exposure as a critical determinant of structural perturbation

Stepping back from His-to-Ala mutations, we instead explored the unique chemical properties of the His triad. Given that histidine residues contain an aromatic ring and can be protonated at physiological pH (p*K*_a_ ∼ 6.5), we considered charge and aromaticity as factors that drive chemical reactivity and structural integrity. We introduced point mutations at each histidine, systematically replacing them with amino acids that reflect one aspect of its chemistry - arginine to mimic the positive charge of a protonated histidine and tyrosine to maintain its aromatic character while shifting its p*K*_a_ to more basic regimes. Six single-point mutations (His15R/Y, His83R/Y, and His87R/Y) were analyzed with respect to WT GM-CSF and one another to identify which chemical property of histidine was most critical to the GM-CSF structure at that position.

We collected ^1^H^15^N HSQC NMR spectra of six GM-CSF variants to quantify CSPs indicative of mutation-induced structural changes. NMR spectra and CD profiles of each GM-CSF variant were well-dispersed and indicative of a folded protein, respectively (**Figure 2A; Supplemental Figure 1**). Mutations at His87 displayed the weakest CSPs (**Figure 2B**), suggesting that the most solvent exposed His87 has minimal impact on the histidine triad chemical environment and GM-CSF structure. Given the evidence that His87 does not appear to strongly impact the GM-CSF structure, we excluded it from further analyses

**Figure 2.**
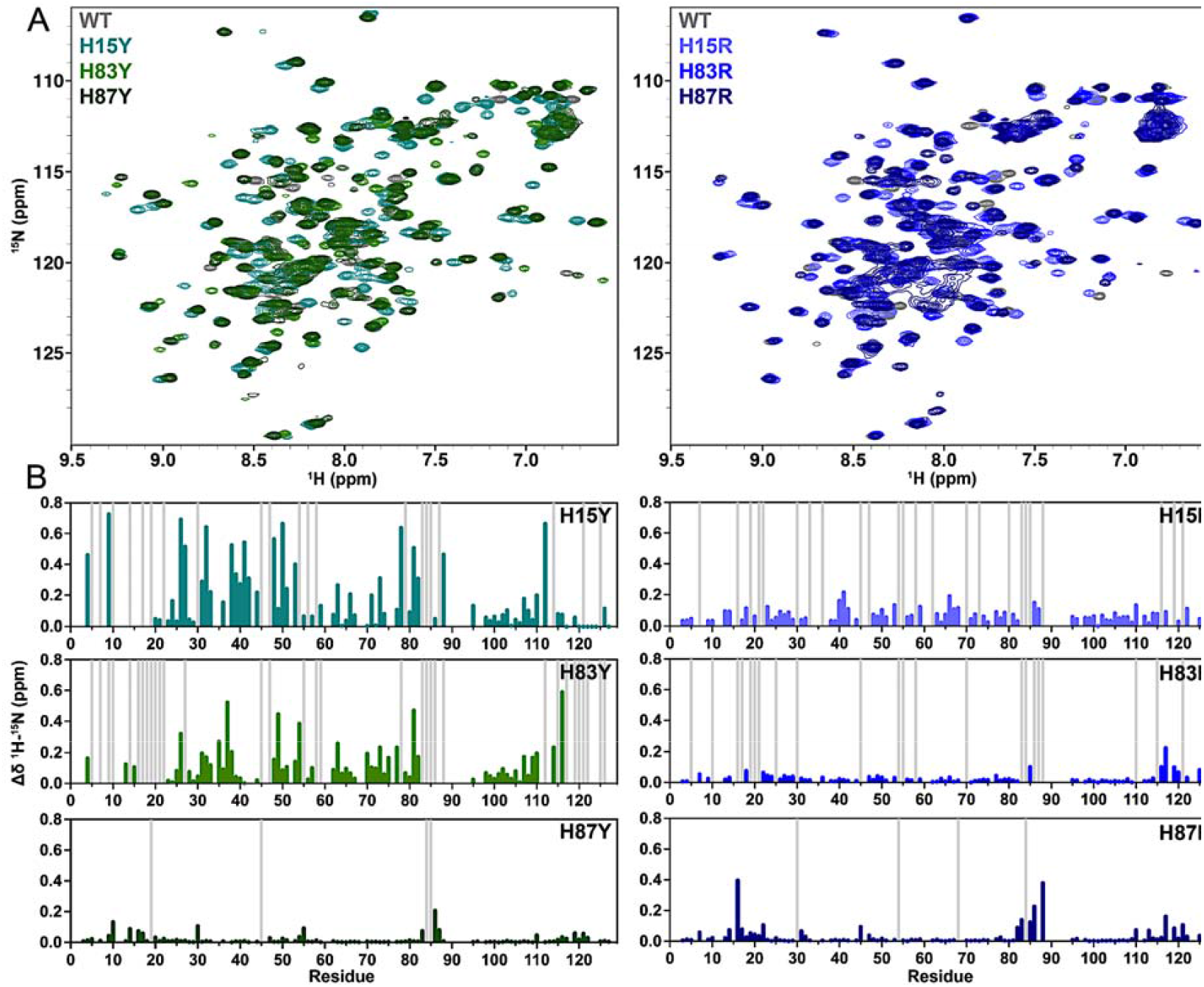
Structural impact of GM-CSF histidine mutations. **(A)** ^1^H^15^N NMR spectral overlays comparing WT GM-CSF (gray) to tyrosine variants (left) and arginine variants (right). **(B)** Per-residue NMR CSPs determined for each GM-CSF variant relative to WT, colored according to the spectra in **(A).** Light gray bars indicate sites of NMR line broadening.

Mutations at His83 caused significant CSPs, with H83Y showing the most pronounced deviations from the WT GM-CSF suggesting that aromaticity alone is not sufficient to maintain the native GM-CSF structure. His15 mutations also induced CSPs, with the Tyr variant of His15, as in H83Y, producing the largest CSPs. However, both H15Y and H15R induce notable line broadening throughout the GM-CSF sequence (**Figure 2B**). Collectively, these data indicate that His15 affects the most significant global structural change beyond the histidine triad site. The overall trend of NMR CSPs reveals that, as histidine residues become increasingly shielded from solvent, mutations disrupt the GM-CSF structure to a greater extent. Most notably, His15 and His83 mutations critically alter the GM-CSF structure, which we predicted would manifest as a change in multi-timescale conformational dynamics.

### NMR relaxation experiments reveal dynamic perturbations in loop regions and N-terminal helices

Building on our insights into structural perturbations from NMR CSPs, we next used *T*_1_, *T*_2,_ and ^1^H-[^15^N] NOE spin relaxation experiments to quantify how the mutations affected the molecular motions of GM-CSF, which are critical to its multifaceted ligand binding interactions. The resulting *R*_1_ and *R*_2_ relaxation rates (**Supplemental Figure 2-5**) shown in **Figure 3** as the *R*_1_*R*_2_ product, account for contributions from anisotropic molecular tumbling and report on *ps–ns* motion with sensitivity to *μs–ms* motion^36^. We visualized the effect of mutations through comparative analysis of GM-CSF variants against WT GM-CSF in correlation plots (**Figure 3B-D**), where deviations from linearity (*ρ*) highlight amino acid-specific differences in flexibility. We find that for the *R*_1_*R*_2_ products, most of the variants show strong dynamic correlation with WT GM-CSF, suggesting negligible change in global tumbling or backbone order (**Figure 3B**). Local changes in flexibility are evident in each variant, particularly in residues contained within *α*1 and *α*4 housing the mutation sites.

**Figure 3.**
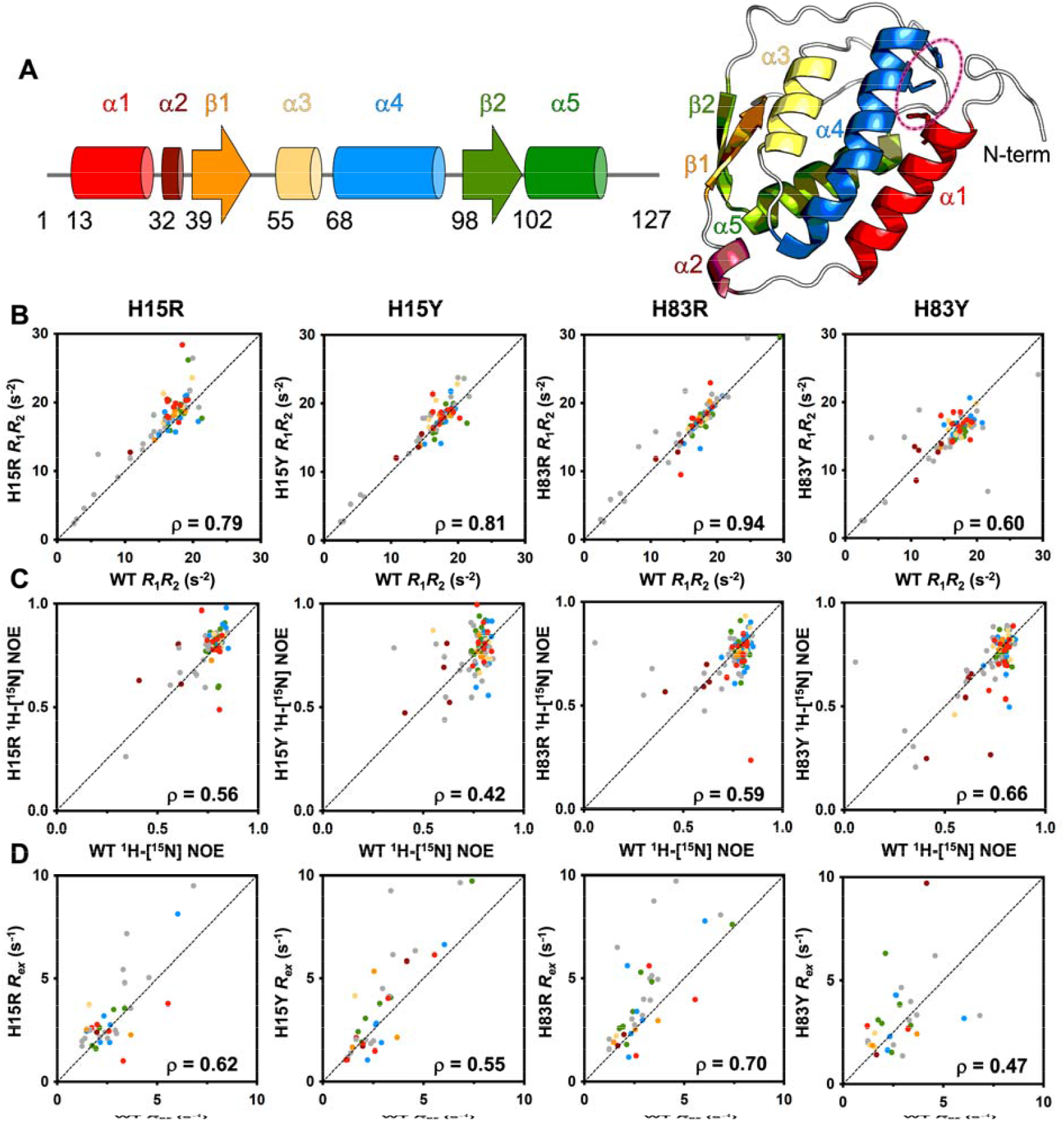
Dynamic effects of GM-CSF mutations. **(A)** Cartoons of the GM-CSF secondary structural elements shown in linear form and mapped onto the PDB: 2GMF. The His triad is circled and data points in subsequent spin relaxation figure panels are colored according to the same structural elements. **(B)** *R*_1_*R*_2_ relaxation parameters plotted as a correlation between WT GM-CSF and each variant. Spearman’s rho values are indicated for each correlation and deviations from linearity highlight specific amino acids with altered local dynamics. **(C)** ^1^H-[^15^N] NOE relaxation parameters plotted as a correlation between WT GM-CSF and each variant, with Spearman’s rho values indicated. **(D)** CPMG-derived *R*_ex_ relaxation parameters plotted as a correlation between WT GM-CSF and each variant, with Spearman’s rho values indicated.

When analyzing ^1^H-[^15^N] NOE values, all variants undergo a change in flexibility compared to WT GM-CSF, in correlation plots and per-residue (**Figure 3C, Supplemental Figure 2-5**). The most prominent effect is generated by the buried His15 residue, where its mutations elevate ^1^H-[^15^N] NOEs across the GM-CSF *α*-helices with no clear pattern, but suppress fast timescale flexibility typically observed in the termini and loops of GM-CSF. Although fast motions appear to be altered in several regions of GM-CSF, these mutation-induced low energy fluctuations do not globally affect the protein and in general, retain many of the characteristics of WT GM-CSF.

We also determined the exchange contribution to transverse relaxation, *R*_ex_, from CPMG relaxation dispersion measurements. These data generally have the poorest correlations with WT GM-CSF, due to significant mutation-induced gain/loss of relaxation dispersion profiles across the sequence. However, H83Y again shows the most substantial deviation in *R*_ex_, as in the *R*_1_*R*_2_ product, demonstrating that perturbation of His83 alters GM-CSF dynamics on the *μs–ms* timescale. In addition to H83Y, *R*_ex_ parameters of H15Y also poorly correlate with WT, suggesting that the charge aspect of the near-neutral His p*K*_a_ (which is removed by a Tyr mutation) is important for preserving WT-like *μs–ms* motions. Collectively, most deviations observed from WT involve residues found in α1 and α4, the N-terminus, and the loop regions of GM-CSF (as seen by the off-diagonal colored dots and structure in **Figure 3**). Both His15 and His83 impact local motion, with His83 most strongly affecting global dynamic parameters.

### Molecular dynamics simulations reveal outward motion of helices in GM-CSF variants

While NMR relaxation experiments provide residue-level detail of the GM-CSF molecular motions, we sought to visualize the atomistic motion using MD simulations. We performed 250 ns MD simulations for WT GM-CSF, H83Y, H83R, H15Y and H15R to assess changes in protein conformational dynamics. Consistent with the relaxation data, we observe subtle fluctuations in the GM-CSF structural ensembles of the variants compared to WT. The conformational flexibility of these mutants have been predicted by prior studies using subsampled AlphaFold2.^37, 38^ Of note, in H83R and H15R *α*1 and *α*4 bend out of plane to sample a greater conformational space (**Figure 4A,B; Supplemental Figure 6**).

**Figure 4.**
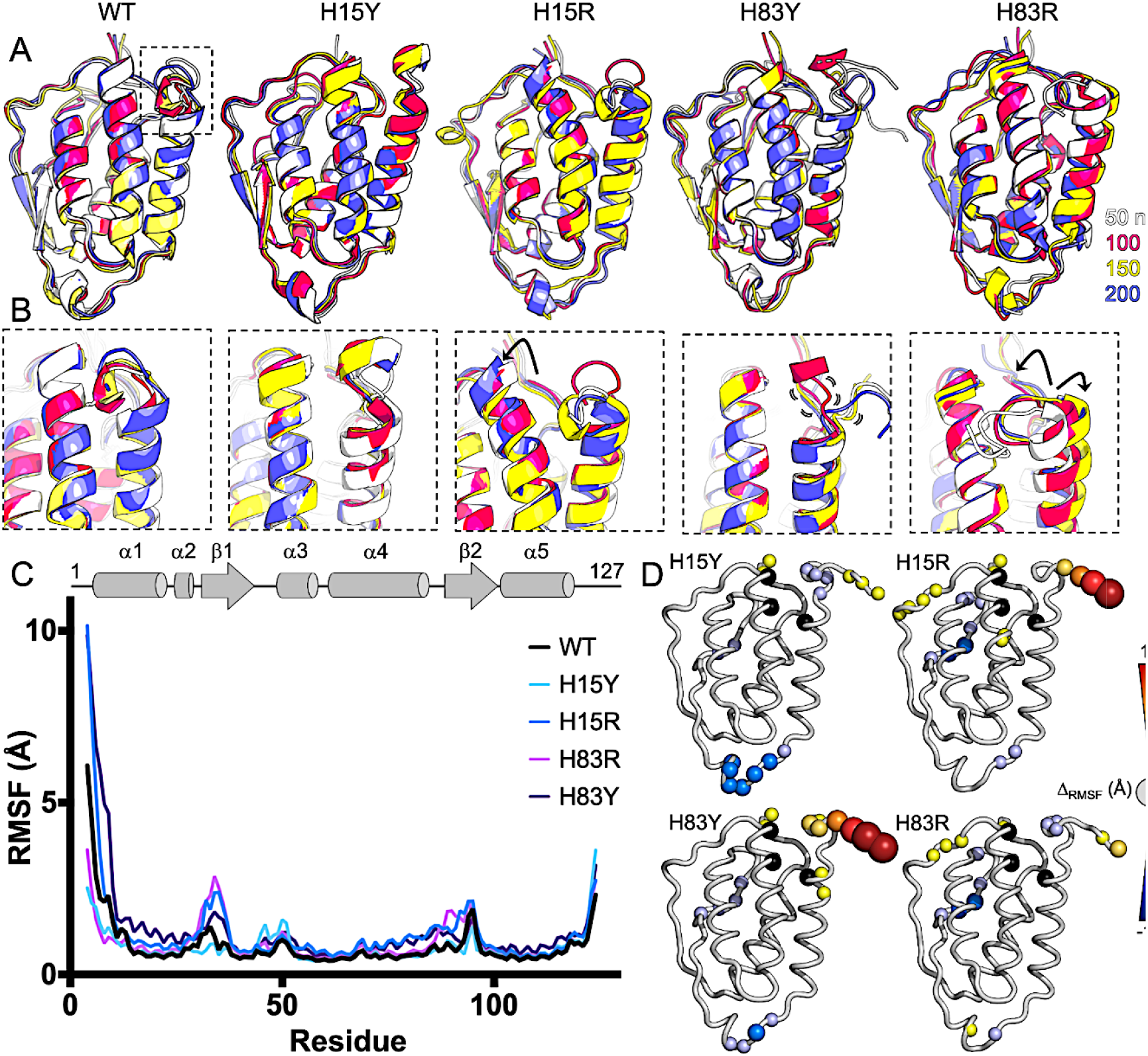
Changes to local GM-CSF conformations. **(A)** Snapshots of WT GM-CSF and variants along the MD trajectories (gray – 50 ns, magenta – 100 ns, yellow – 150 ns, blue – 200 ns). **(B)** The same MD ensembles focused on *α*1 and *α*4 containing the His triad. Notable conformational changes (H15R, H83R) are indicated by arrows and subtle molecular motions are also indicated for H83Y. **(C)** Per-residue RMSF derived from MD simulations of WT and GM-CSF variants. Traces are colored according to the legend and the secondary structure of GM-CSF is shown as a cartoon at top. **(D)** Summary of mutation-induced changes in RMSF (Δ_RMSF_ (mutant – WT)) mapped onto the GM-CSF structure. Sphere sizes correlate to the magnitude of change. Warm colors highlight greater flexibility of GM-CSF variants and cool colors highlight greater flexibility in WT GM-CSF.

To quantify these differences, we calculated the root mean square fluctuations (RMSF) of the protein backbone for all variants and WT GM-CSF (**Figure 4C**). We find that the N-terminus experiences higher fluctuation in both H15R and H83Y relative to the WT, while H15Y attenuates backbone fluctuations. Notably, the RMSF profile of H83R most resembles that of the WT (**Figure 4D**), which in light of its NMR relaxation parameters, suggests that electrostatic effects, rather than aromaticity, drives the dynamic impact of this residue. In His15, on the other hand, both tyrosine and arginine mutants alter the intrinsic motions of WT GM-CSF, indicating that both aromaticity and charge influence protein dynamics at His15 under neutral pH conditions.

### Histidine mutations do not alter GM-CSF ligand binding propensity

Building on the subtle structural fluctuations seen by MD simulations, we examined the impact of histidine mutations on physiologically relevant ligand binding. We tested the mutations with the highest CSPs from WT, namely H15Y and H83Y, with a larger biological molecule, heparin, previously shown to have pH-dependent binding to WT GM-CSF^14, 15, 35^, as well as a the ATP molecule, which was shown to have enhanced production through a GM-CSF signaling pathway^39, 40^. Both ligands carry significant negative charges with an overall negative charge consistent with the larger heparin molecule tested. We collected ^1^H-^15^N HSQC titrations with increasing concentrations of dp6 heparin. Subsequent CSPs were stronger for the H15Y and H83Y variants than WT (**Figure 5A**). We also quantified the apparent binding affinity (*K*_d_) from CSP trajectories and, as expected, found weak affinities of H83Y (0.9 mM) and H15Y (0.8 mM) for heparin, though tighter than the 1.2 mM of WT GM-CSF (**Supplemental Figure 7**). To model the binding, we used the ClusPro server^41, 42^ with a heparin-specific algorithm optimized for shallow charged pockets. The larger heparin molecule binds to basic patches on the exterior of GM-CSF, both adjacent to the His triad and along the *α*1 face, with minimal difference between the variants (**Figure 5B**). Thus, slight differences in binding of heparin likely occur from structural changes to GM-CSF caused by the histidine mutations, but not solely within the pocket itself.

**Figure 5.**
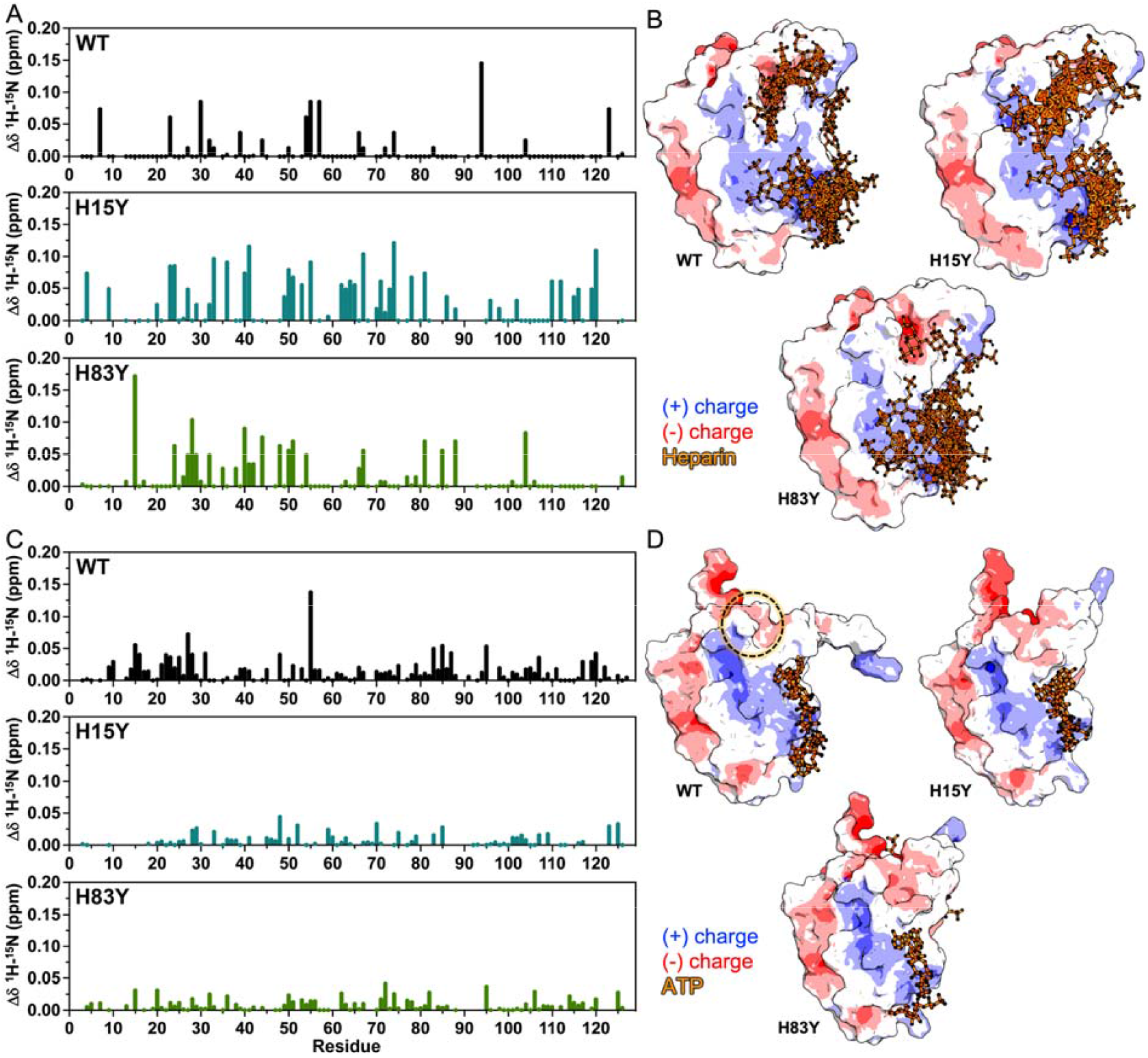
Interactions of GM-CSF with ligands. **(A)** ^1^H-^15^N NMR CSPs caused by 1.8 mM dp6 heparin binding to WT (black), H15Y (cyan), and H83Y (green) GM-CSF. **(B)** Binding poses of hepari s with the same GM-CSF proteins predicted by ClusPro. GM-CSF structures are colored according to surface charge, and docked heparins are shown in orange. **(C)** ^1^H-^15^N NMR CSPs caused by 1.1 mM ATP binding to WT (black), H15Y (cyan), and H83Y (green) GM-CSF. **(D)** Binding poses of ATP ligands with the same GM-CSF proteins predicted by AlphaFold3. GM-CSF structures are colored according to surface charge and docked ATP molecules are shown in orange. The His triad is indicated by the dashed circle.

We repeated titrations with ATP, a much smaller molecule than dp6 heparin which potentially could bind within the histidine pocket. NMR CSPs were smaller in magnitude, though saturation of GM-CSF occurred at ATP concentrations lower than required for dp6 heparin. In contrast to heparin, ATP induced the largest CSPs in WT GM-CSF (**Figure 5C**) and the apparent *K*_d_ followed a trend with WT GM-CSF having a slightly tighter affinity (0.35 mM) than H15Y (0.45 mM) and H83Y (0.47 mM), though still a weak interaction. The greater affinity of GM-CSF for ATP could indicate some direct binding within the histidine pocket that is disrupted by the tyrosine mutations, rather than the diffuse interactions of heparin. The effects of a tyrosine mutation, where tyrosine destabilizes the helix such that the histidine pocket is more accessible, are counterbalanced by the removal of a potential charged interaction between GM-CSF and the ligand. We then used AlphaFold3 to model the GM-CSF-ATP complex (**Figure 5D**), which showed ATP sampling multiple binding sites on the positive face of the protein, but none in the histidine pocket.

Since neither algorithm predicts the histidine triad as a dominant binding sites for either ligand, we investigated whether a change in the distance between the *α*1 and *α*4 helices corresponded to the minor changes in binding affinity. We found that at no point during these MD simulations was the interior of GM-CSF core accessible to solvent and that the solvent accessible surface area does not significantly change between the WT and GM-CSF variants (**Supplemental Figure 8, 9**). The seemingly accessible channel is lined with hydrophobic residues that hinder the ability for solvated ligands to bind beyond the shallow pocket of the triad.

In conjunction with the NMR, these data suggests that at neutral pH, the histidine triad, even when manipulated to favor a more flexible state, does not drive ligand binding. Rather, these interactions seem to be transiently mediated along the positive electrostatic surface of the protein.

## Discussion

The GM-CSF histidine triad is an understudied region of its structure. The compact cytokine fold of GM-CSF renders it difficult to discern how multiple non-overlapping functions could be contained within its scaffold. The clustering of three His residues in GM-CSF has been demonstrated in prior work to be a pH sensor for ligand binding^14, 15^, and here, to be a critical stabilizing force for the native fold. Using NMR spectroscopy and computational tools, we showed that GM-CSF is sensitive to mutations at these key histidine sites at neutral pH. Conventional alanine substitutions strongly destabilized the protein, underscoring the importance of amino acid substitutes that replicate the distinct chemical properties of histidine (aromatic ring, positive charge).

From the analysis of six single-point mutations, the results can be categorized along two axes: solvent exposure and chemical characteristics. Specifically,, the structural and dynamic changes in GM-CSF, assessed through NMR CSPs, spin relaxation, and heteronuclear NOE, are largely influenced by the solvent exposure or burial of the mutated residue. Notably, mutations at His15, which is the least solvent exposed, caused the most significant alterations to the structure and dynamics of GM-CSF when its charge or aromaticity was altered. Substitutions of different amino acids at the same residue position yielded distinct NMR spectral patterns, reinforcing that mutational effects in GM-CSF are highly position- and type-dependent and not uniform across the samples.

Following these structural and dynamic perturbations, we anticipated that mutations causing the helices to move away from each other would enhance ligand binding similarly to acidic pH. Consequently, we titrated both heparin and ATP, known ligands of GM-CSF, to quantify binding affinity. We were surprised that at neutral pH, none of the variants displayed a significant change in ligand affinity, suggesting that any one histidine alone is not sufficient to enhance binding.

As a result, we asked whether there was a correlation between the “openness” of the His pocket and any changes in binding affinity. These data reveal that even though mutations caused notable structural and dynamic perturbations near the GM-CSF N-terminus, they alone are not sufficient to facilitate binding. Instead, at neutral pH, the interactions of GM-CSF with its ligands seem to be primarily driven by electrostatic interactions via distinct positively and negatively charged faces of the protein. This may explain why neither NMR experiments nor computational predictions show altered binding interactions in these sites at neutral pH. Such transient interactions driven by electrostatics have been documented in many other proteins^43, 44^.

These results provide a more comprehensive model of how GM-CSF may toggle between its different functions: at neutral pH, the electrostatic, “sticky” surface drives transient, low-affinity interactions with its ligands. At acidic pH, which physiologically may correlate to inflammatory compartments^45^, the ionized histidine binding pocket may aid in increasing the specificity of molecular interactions. Here, mutation of specific His residues reveal the conformational fluctuations that underscore the plasticity of this site.

## Acknowledgments

This work was supported by NIH grant R01 GM144451 (to GPL).

