## Supporting Information for "Exploring the GM-CSF Histidine Triad as a Modulator of Structure, Molecular Motion, and Ligand Binding"

**Figure S1.** CD spectra of WT GM-CSF and variants

**Figure S2-S5.** Relaxation parameters for WT GM-CSF and variants

**Figure S6.** Distance between residue 15 and 83 in MD Simulations.

**Figure S7.** ^1^H^15^N HSQC NMR spectra and *K*_d_ curves of WT, H15Y, and H83Y with heparin (dp6) and ATP

**Figure S8.** Representation of GM-CSF with hydrophobic and aromatic residues highlighted.

**Figure S9.** Solvent accessible surface area changes over time.

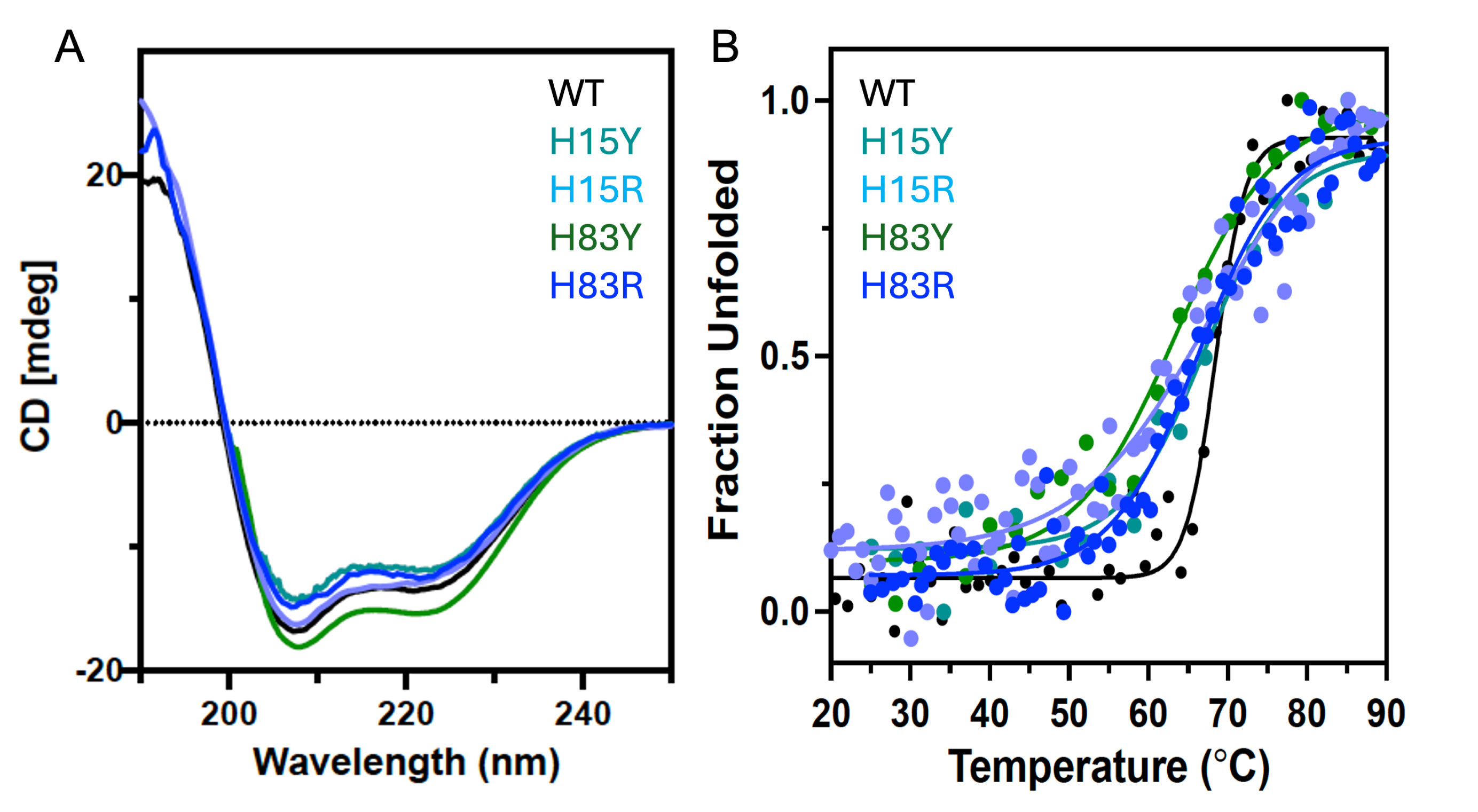

**Figure S1.** **(A)** CD spectra of WT (black), H15Y (cyan), H15R (light blue), H83Y (green), and H83R (blue) GM-CSF. Secondary structure profiles are highly similar. **(B)** Thermal denaturation experiments monitoring CD ellipticity at 220 nm for the same GM-CSF samples.

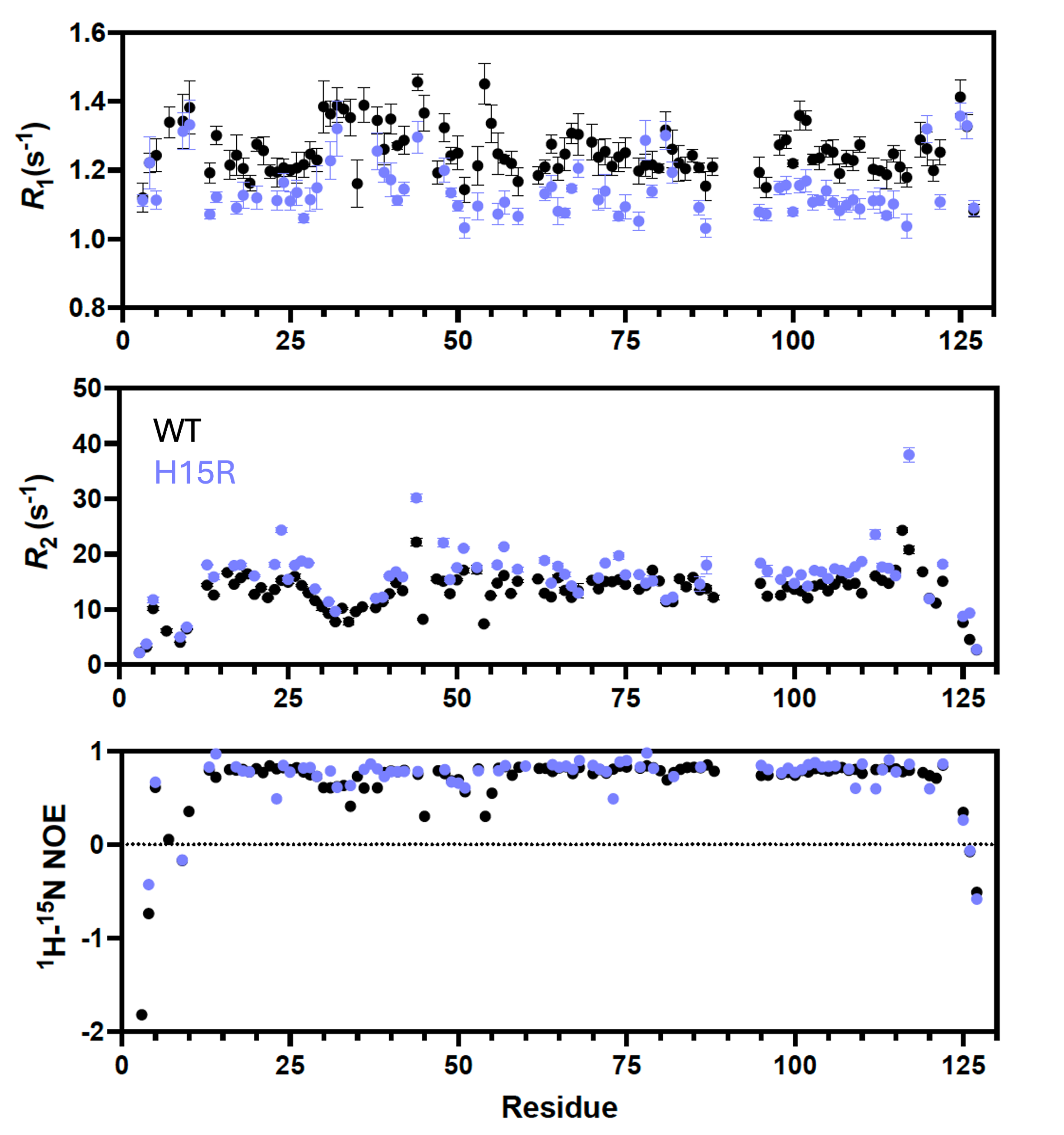

**Figure S2**. Per-residue *R*_1_, *R*_2_, and ^1^H-[^15^N] NOE relaxation parameters for WT GM-CSF (black) and H15R GM-CSF (light purple).

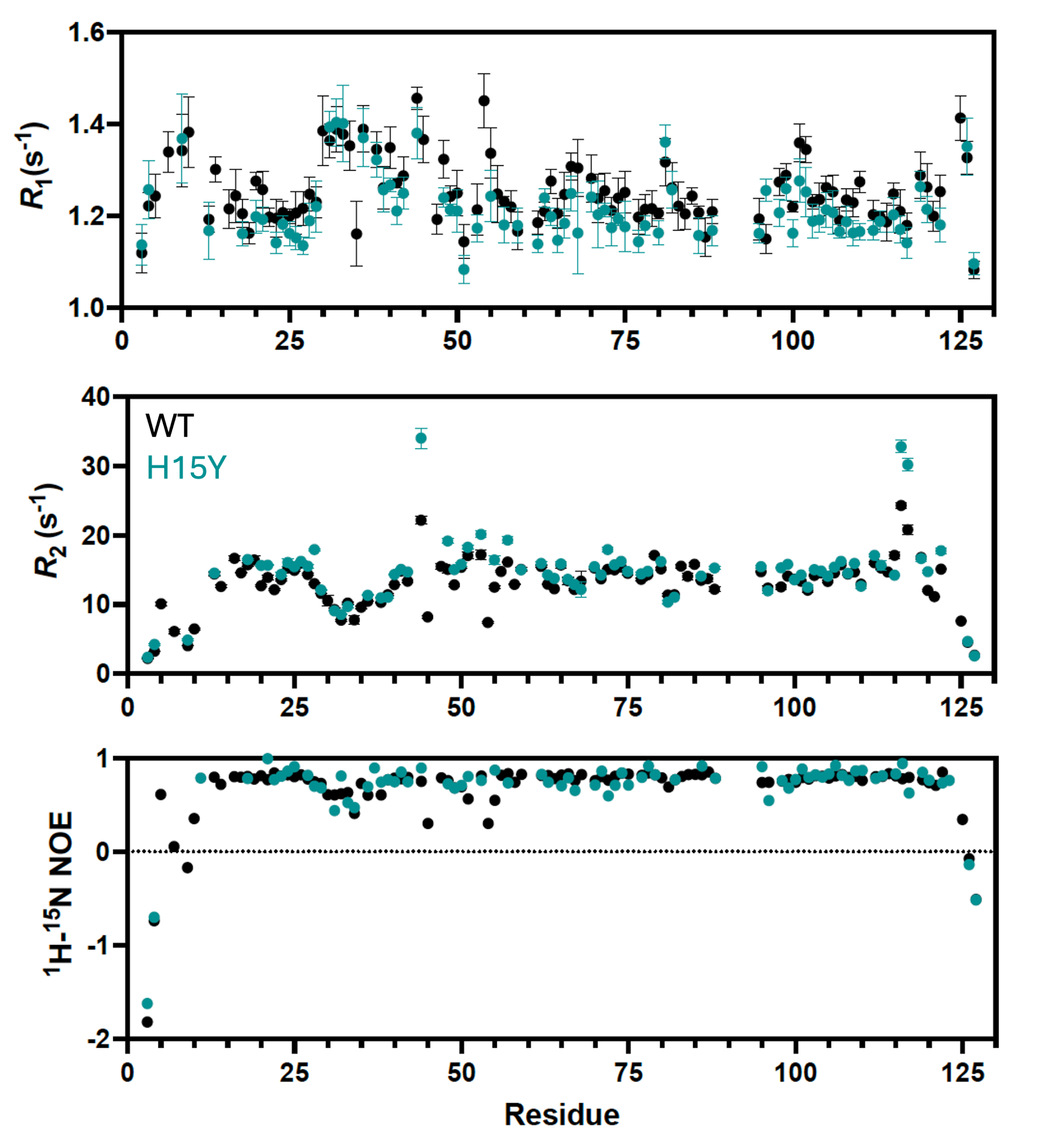

**Figure S3**. Per-residue *R*_1_, *R*_2_, and ^1^H-[^15^N] NOE relaxation parameters for WT GM-CSF (black) and H15Y GM-CSF (cyan).

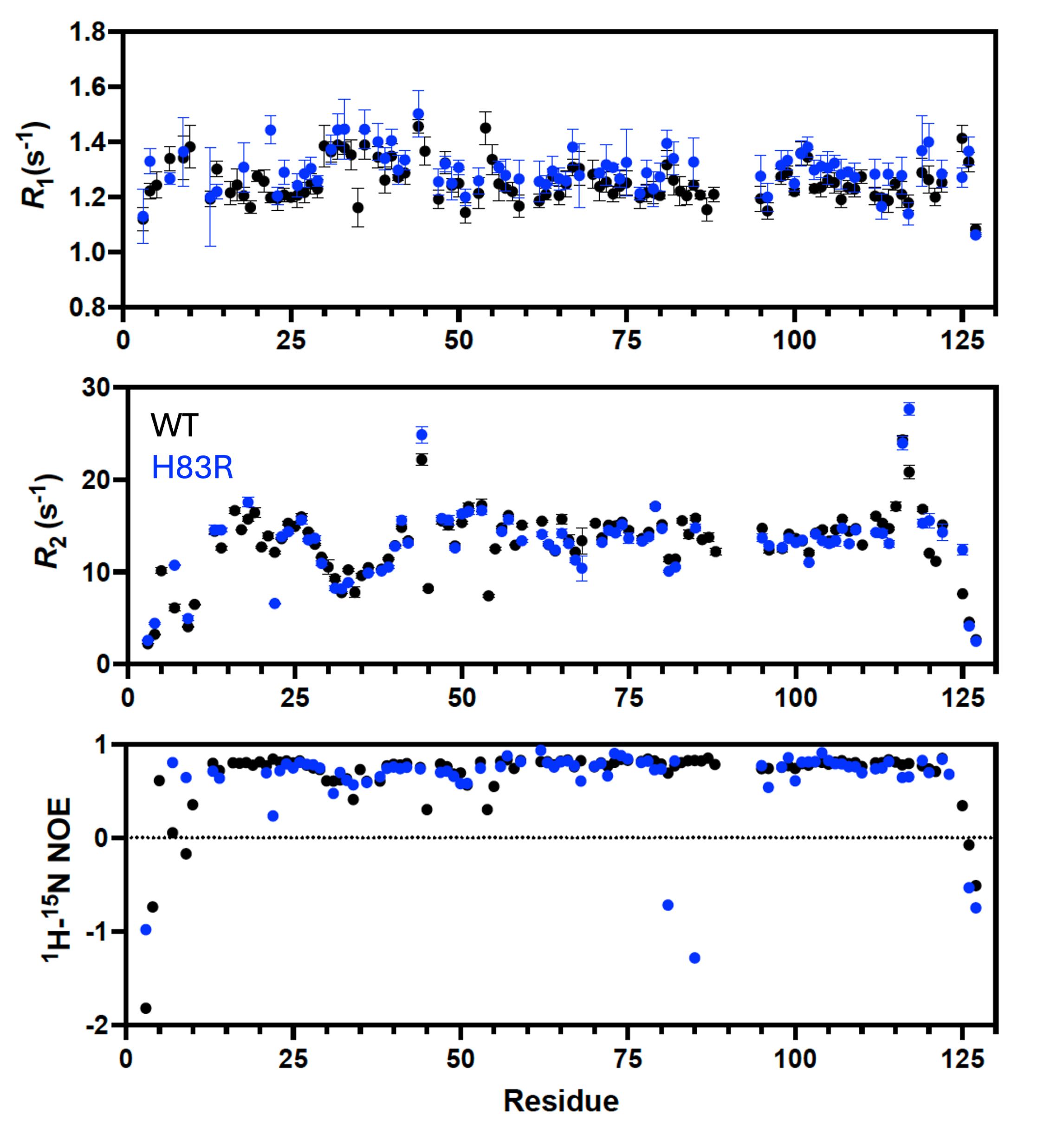

**Figure S4**. Per-residue *R*_1_, *R*_2_, and ^1^H-[^15^N] NOE relaxation parameters for WT GM-CSF (black) and H83R GM-CSF (blue).

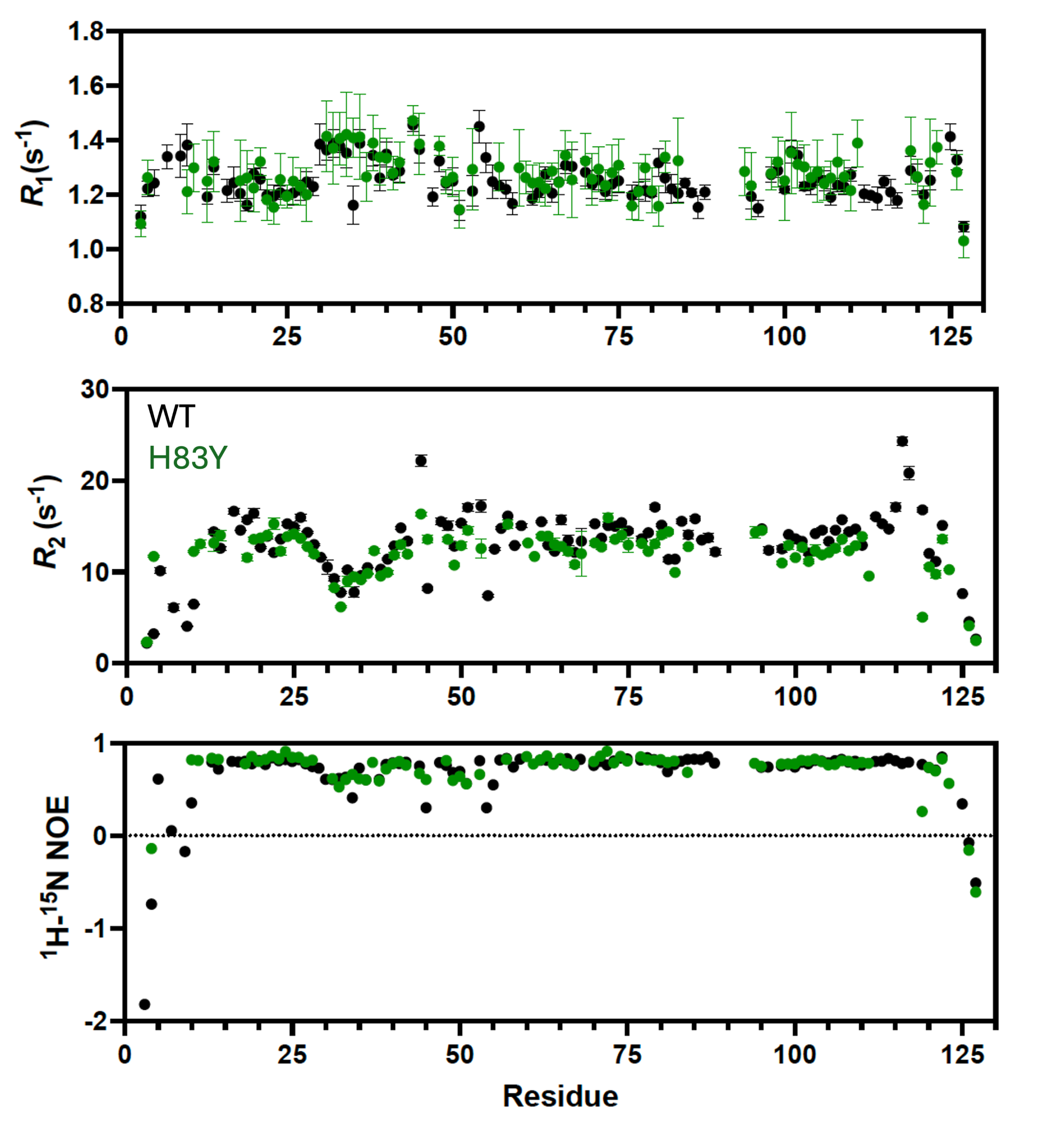

**Figure S5**. Per-residue *R*_1_, *R*_2_, and ^1^H-[^15^N] NOE relaxation parameters for WT GM-CSF (black) and H83Y GM-CSF (green).

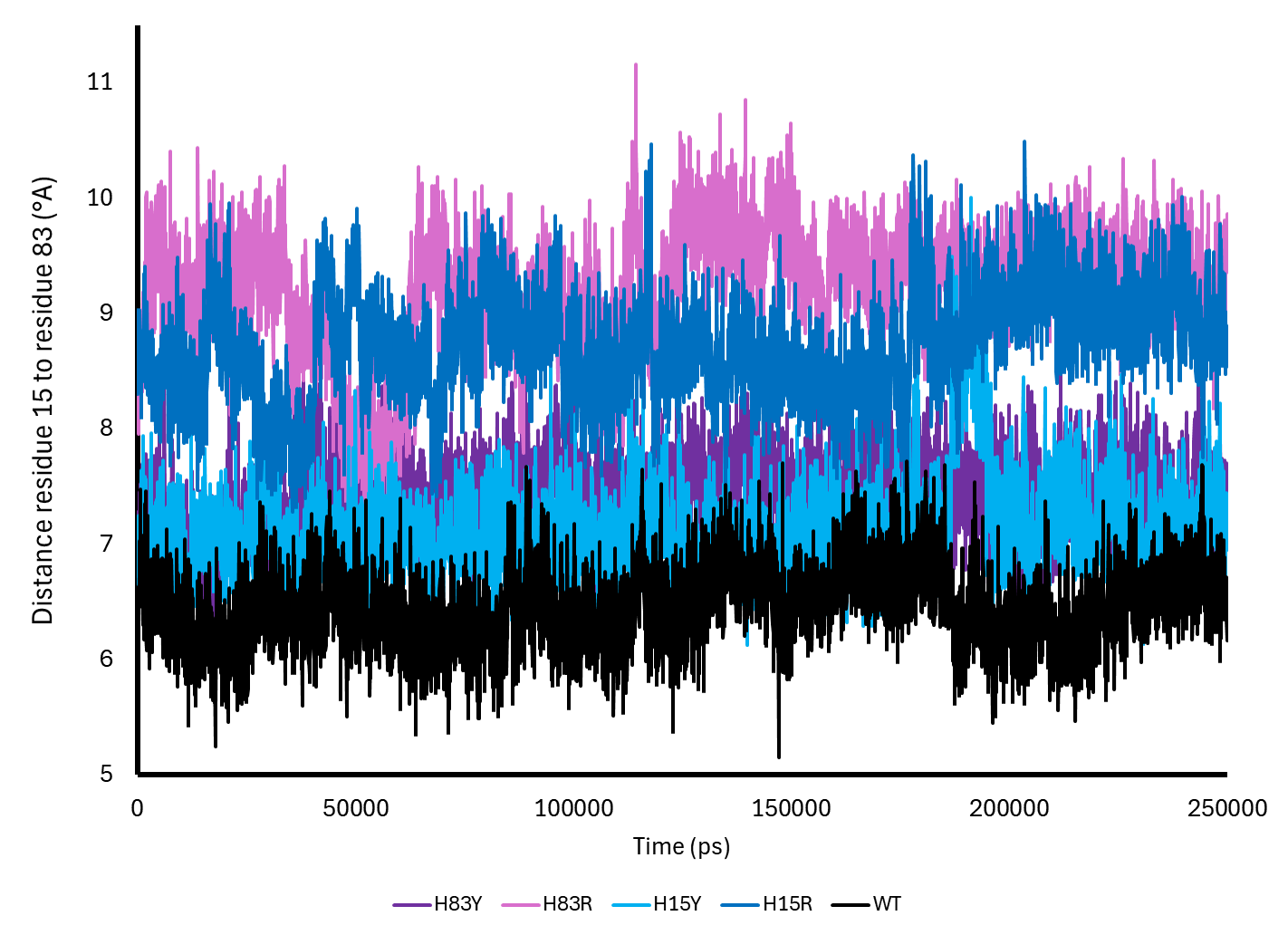

**Figure S6** Distance between residue 15 and 83 in MD simulations over time.

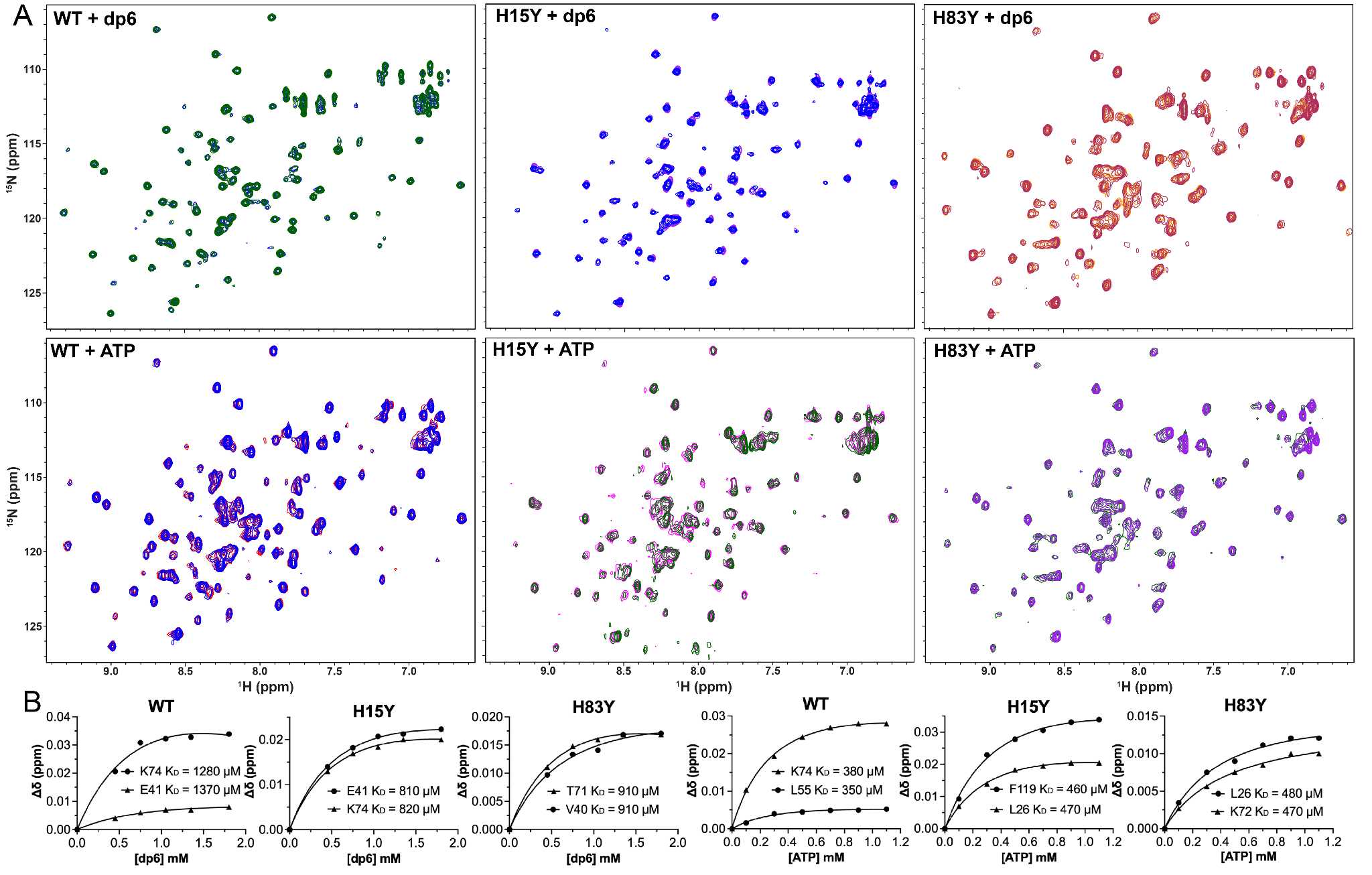

**Figure S7.** ^1^H^15^N HSQC NMR spectra and apparent *K*_d_ curves for WT, H15Y, and H83Y GM-CSF with heparin (dp6) and ATP. **(A)** ^1^H^15^N HSQC spectra overlay of WT, H15Y, and H83Y GM-CSF with 1.8 mM dp6 and 1.1 mM ATP. **(B)** *K*_d_ curve fits of ligand titration from representative peaks of each variant.

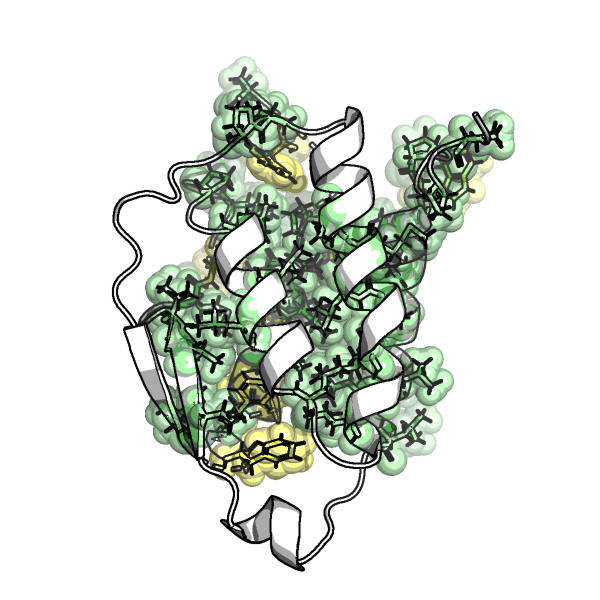

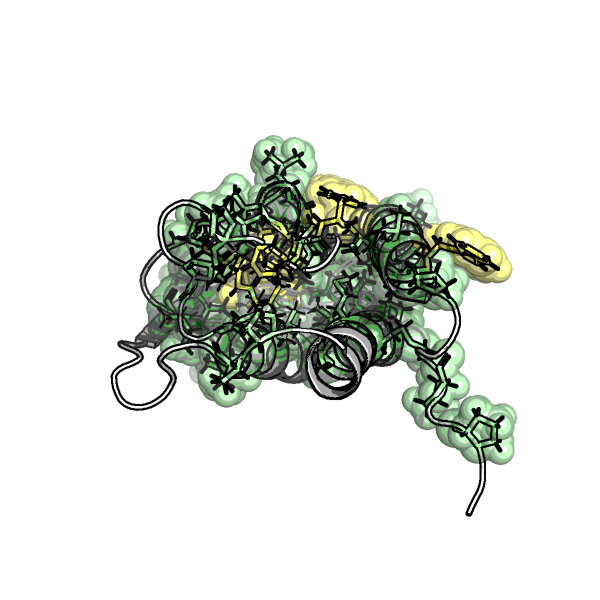

**Figure S8. Representation of GM-CSF with hydrophobic and aromatic residues highlighted.** Hydrophobic residues in GM-CSF pocket, shown in green, while aromatic residues are shown in yellow.

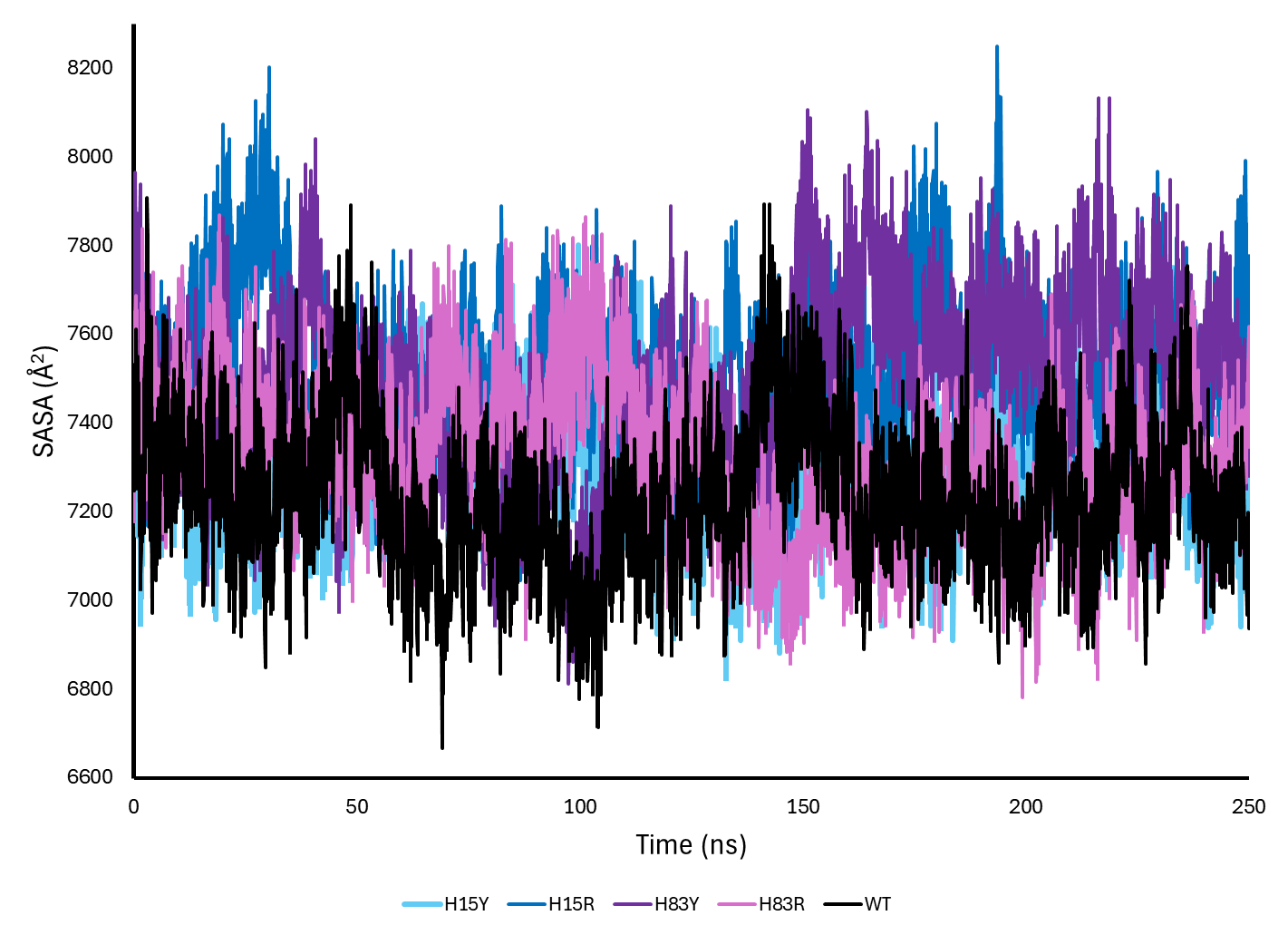

**Figure S9.** Solvent accessible surface area (SASA) changes over time for GM-CSF variants in MD simulations.
